# The variability of expression of many genes and most functional pathways is observed to increase with age in brain transcriptome data

**DOI:** 10.1101/526491

**Authors:** Veronika R. Kedlian, Handan Melike Donertas, Janet M. Thornton

## Abstract

Ageing is broadly defined as a time-dependent progressive decline in the functional and physiological integrity of organisms. Previous studies and evolutionary theories of ageing suggest that ageing is not a programmed process but reflects dynamic stochastic events. In this study, we test whether transcriptional noise shows an increase with age, which would be expected from stochastic theories. Using human brain transcriptome dataset, we analysed the heterogeneity in the transcriptome for individual genes and functional pathways, employing different analysis methods and pre-processing steps. We show that unlike expression level changes, changes in heterogeneity are highly dependent on the methodology and the underlying assumptions. Although the particular set of genes that can be characterized as differentially variable is highly dependent on the methods, we observe a consistent increase in heterogeneity at every level, independent of the method. In particular, we demonstrate a weak but reproducible transcriptome-wide shift towards an increase in heterogeneity, with twice as many genes significantly increasing as opposed to decreasing their heterogeneity. Furthermore, this pattern of increasing heterogeneity is not specific but is associated with a wide range of pathways.

## INTRODUCTION

Ageing is commonly defined as a time-dependent decrease in the functional and structural integrity of an organism. Despite the ubiquity of ageing in all living organisms, the molecular mechanisms responsible still require further elucidation. According to recent studies, ageing differs phenotypically among individuals, including monozygotic twins [1,2] and within tissues from the same individuals [3]. Researchers have observed an age-related increase in variability in the epigenome [4,5] and transcriptome [6] of genetically identical samples, which may underlie the phenotypic differences. Age-related expression variability has been detected in many different cell and tissue types including mice stem cells, cardiomyocytes and immune cells [7–9], rat neural retina [10], fruit-fly, mice and human brain [6,11–14] as well as human pancreas, lung, blood, skin, fat and human fibroblasts *in vitro* [13,15–17]. Despite these reports, there is no agreement on the underlying mechanisms, extent and functional consequences. Suggested mechanisms include somatic [7,15] and germline mutations [11,17], changes in the DNA methylation [9,17,18] and chromatin modifications [5] and resulting chromatin compaction [12] as well as global dysregulation, caused by the change in transcription factor or miRNA expression [19].

Both genome-wide and hypothesis-driven approaches have been employed to explore the extent of expression variability with age. Among the former, some show a transcriptome-wide increase [6,9,12,13,15], while others focus only on those genes showing significant changes in their variability. Brinkmeyer-Langford *et al.* [11] reports that an equal number of genes significantly increase or decrease their expression variability, whereas a recent study from Vinuela *et al.* [17] shows more genes decreasing rather than increasing their expression variability [17]. Hypothesis-driven studies mostly show an increase in variability for the genes measured [7,8,16], whereas Warren *et al.* [20] suggests this might be specific only to the non-renewing tissues. Similarly, Ximerakis *et al*. [14] shows that change in transcription variability is in different directions in different cell types of mouse brain. The reports also vary in terms of the functional association of this variability. While some consider that increase in variability is widespread [6,12], others report that variability is concentrated in various cellular functions [10,11,18,21] – although these functions also differ between reports.

Age-dependent change in the expression variability is difficult to address due to the inherent noise in expression and the influence of other factors on variability. Thus, the data pre-processing steps to disentangle variability from the biological and technical confounders is of importance. Another technical aspect is the method to measure the change in the variability. Most studies tested for age-related change in the expression variability using either *grouped* (Bartlett’s test, Levene’s test, permutation test)[7,11,20] or *regression-based* tests (linear and loess regression)[6,10,17,18], with a few others using *correlation-based* approaches (gene co-expression, intra-class correlations)[21,22]. However, to our best knowledge, the effects of different batch-correction strategies and different methods to measure variability have not been explored on the same data.

In this study, we undertook a comprehensive investigation of the ageing-related change in expression variability, using human brain expression dataset. We employed different pre-processing and variability measures and analysed transcriptome-wide and gene-level changes in gene expression variability and the associated functions.

## RESULTS

In order to study the change in gene expression variability during ageing, we used one of the biggest published human brain transcriptome datasets, generated using microarray technology [23]. We limited the age range to between 20 and 80 years (Figure 1A), resulting in RNA expression data for 147 prefrontal cortex samples. We excluded prenatal, infant and childhood samples (up to 20 years old), because their expression levels will be inherently coupled to developmental processes in the brain. We applied four batch correction strategies to account for technical and biological confounders (Supplemental Figure 1): i) only quantile normalization (QN), ii) QN followed by linear regression (regression), iii) QN followed by ComBat [24], and iv) QN followed by Surrogate Variable Analysis (SVA) [25]. Regression and ComBat are supervised approaches, i.e. known covariates should be supplied to the algorithm, whereas SVA estimates covariates from the data. We provide the results from Regression and SVA in the main text to include one supervised and one unsupervised approach. The results from other correction strategies are given in Supplemental Data in comparison with the Regression and SVA approaches (Supplemental Figures 2-8).

**Figure 1.**
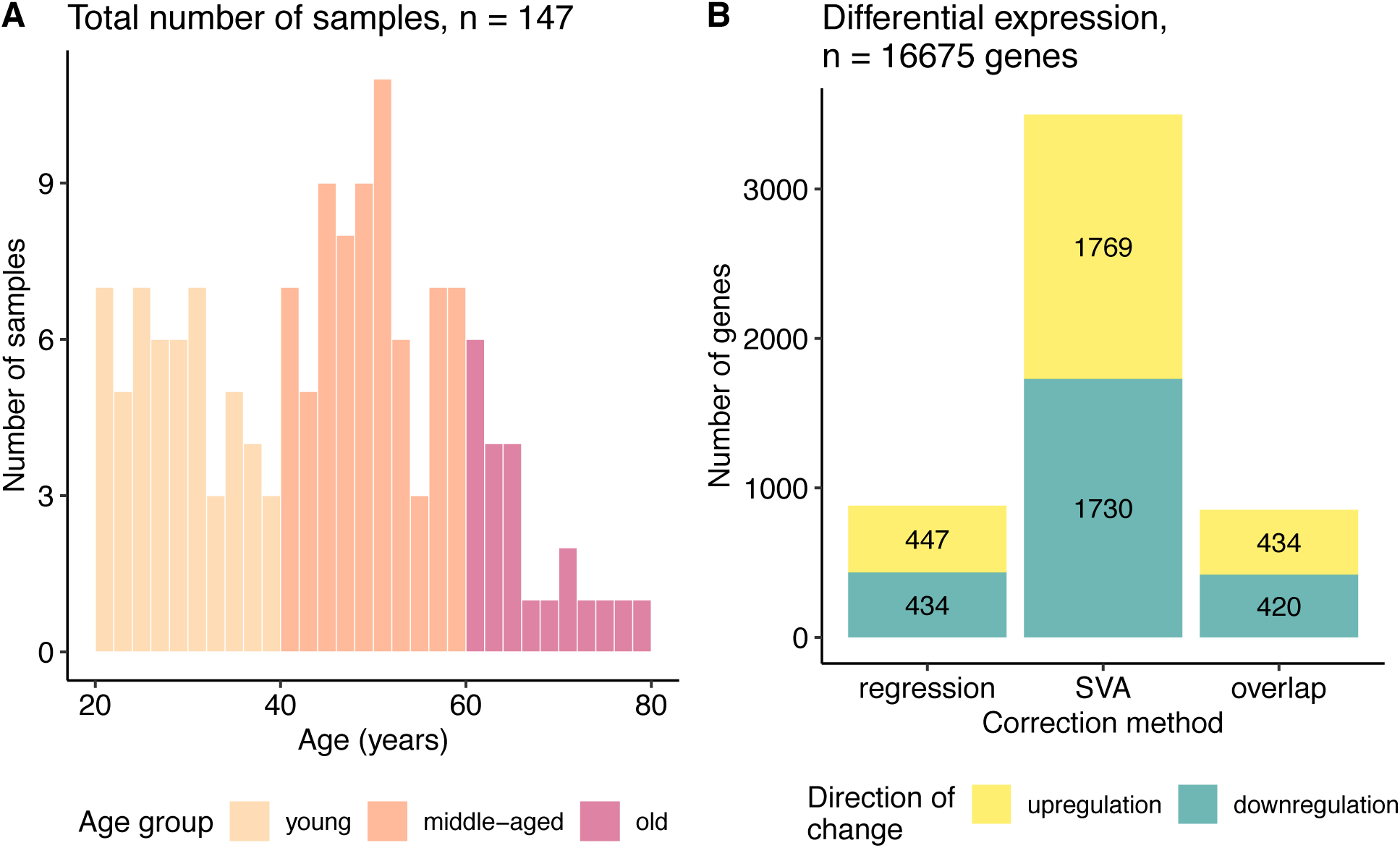
Data characterization. (**A**) Age distribution of the samples used in the study. (**B**) Bar plot of the number of genes differentially expressed with ageing identified after regression and SVA correction and their overlap. The colour represents direction of change: yellow – genes upregulated and blue – downregulated with age.

### Analysis of the differentially expressed genes

First, we defined differentially expressed (DE) genes, based on the significance of the regression coefficients (FDR corrected p <= 0.05) for the linear model using the gene expression values as the dependent and age as the independent variable (see Methods). By applying two different pre-processing approaches on the data prior to DE estimation the number of DE genes was very different. SVA correction yielded 3499 DE genes, compared with 881 DE genes found by regression (Figure 1B, Supplemental Table 1). Nevertheless, 854 genes overlapped between the SVA and regression results, which constituted a quarter of genes found after SVA and 96% of the genes identified after regression correction. Quite high overlap was consistent with the strong correlation between the expression level changes for the regression and SVA corrected data (Spearman ρ = 0.85, Supplemental Figure 3A).

To explore the biological processes affected by these changes in gene expression, gene set enrichment analysis was performed separately on the regression and SVA corrected data (Supplemental Table 2). It revealed 125 (out of 191 (regression) and 160 (SVA) categories) shared Gene Ontology Biological Process (GO BP) categories that were downregulated in the ageing brain. Cognitive-function related GO terms, such as modulation of synaptic transmission, learning or memory, constituted a substantial fraction of these GO terms. In contrast, the number of the upregulated GO terms was much smaller and only 8 GO terms overlapped (out of 22 (regression) and 12 (SVA) categories) between the correction approaches, including detoxification, stress response to metal ions and cilium organization GO categories (see Supplemental Table 2).

### Analysis of the differentially variable genes

Two different strategies were employed to measure change in the gene expression variability with age, namely continuous and grouped approaches. *The continuous approach* detects continuous monotonic change in variation from 20 to 80 years of age. *The grouped approach* compares the gene expression variation between two age groups: young (20 – 40 years old, N = 53) and old (60 – 80 years old, N = 22). Figure 2 illustrates the principles of these approaches and shows that the change in variability can be combined with any dynamics in the mean gene expression (upregulation, downregulation, no change). We checked if the changes in gene expression variability is confounded by the changes in gene expression level, but did not observe any relationship (Supplemental Figure 10, Fisher’s test p = 0.11, Odds ratio = 1.05).

**Figure 2.**
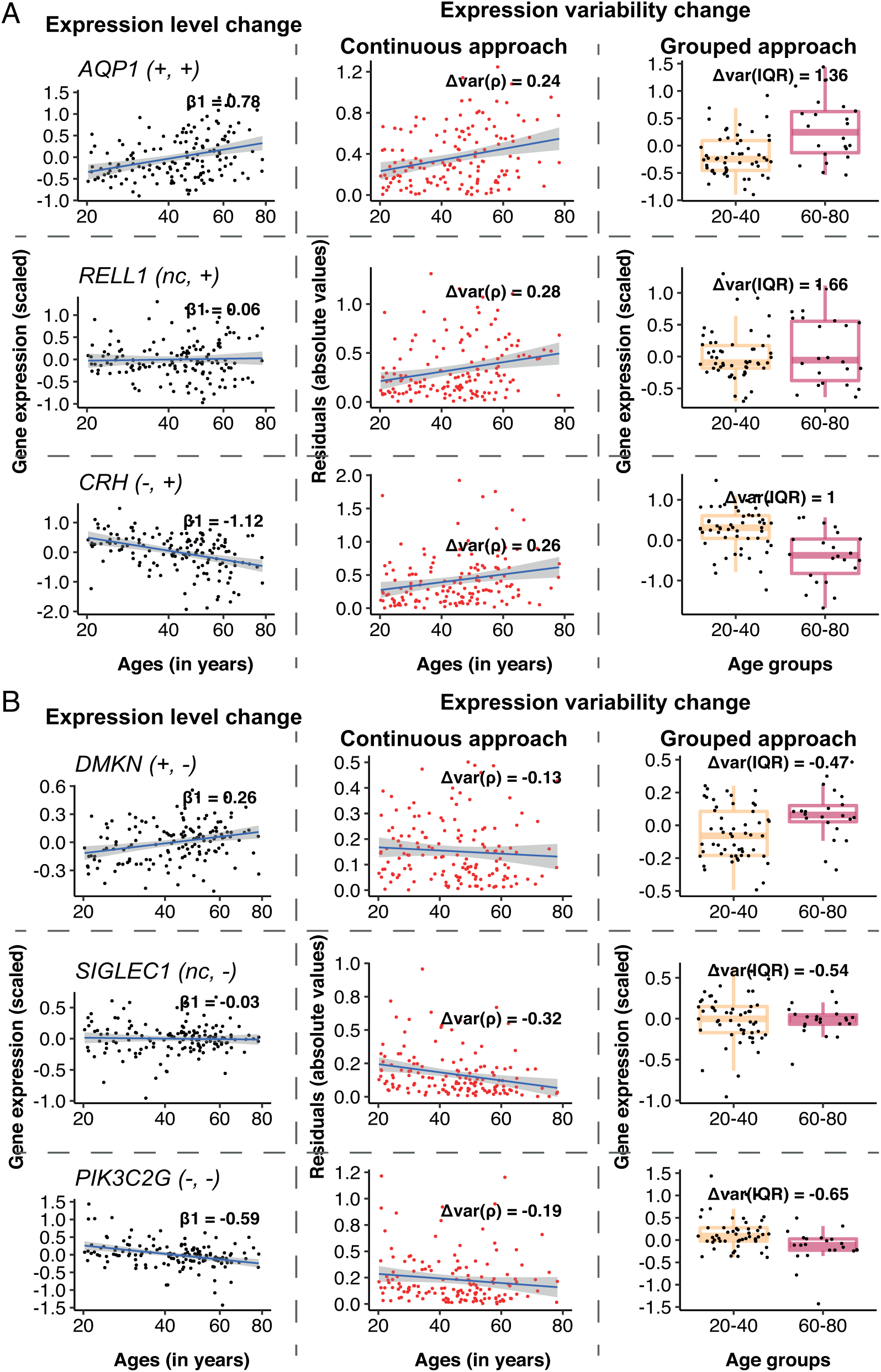
Changes in gene expression and its variability with age for some individual genes, using the different approaches. Example genes are chosen that increase (**A**), or decrease (**B**) expression variability with age, when the mean gene expression either increases, does not change or decreases. The types of change, for expression and variability respectively, is shown in the parenthesis following the gene name, for each row. Genes were selected to have the biggest absolute values of Δvar(ρ) and Δvar(IQR) as well as demonstrate significant increase, decrease or no change in the expression level with age. The first column to the left illustrates mean expression level (regression-corrected) plotted against individual’s age on the x^0.25^ – transformed scale. The regression line is colored in blue, with the β1 coefficient from the linear regression shown on the graph. The middle column illustrates the continuous approach to measure differential variability. Absolute values of the residuals (in red) from the regression line are plotted against age and the regression line between residuals and age (in blue) is drawn for illustrative purposes. The Spearman correlation estimates, Δvar(ρ) between the residuals and age are displayed on the graph and used in the subsequent analysis. The last column on the right illustrates the grouped approach to calculate differential variability. Gene expression levels (regression-corrected) of the individuals from the “young” (20 – 40 years old) and “old” (60 – 80 years old) groups are represented in the corresponding boxplots. A small random deviation (jitter) from the x-axis is applied for better visualization. Δvar(IQR), the fractional change in the variability in the “old” group, as compared to the “young”, is displayed on the graph.

*In the continuous approach*, we first fit a linear model to explain age-dependent change in expression (Figure 2, first column) and then used the residuals from this model to represent the variability. To measure change in the expression variability with age, we calculated the Spearman correlation coefficient (Δvar(ρ)) between the absolute value of residuals and age (Figure 2, middle column). The Δvar(ρ) measures ranged between −0.32 and 0.36 and were normally distributed (Shapiro-Wilk test, p > 0.05, see Methods) (Figure 3A). The distributions were moderately, but significantly (Wilcoxon test, p < 2.2e-16), shifted towards positive values for both correction methods. Although the shift in the distribution was small, 57% to 63% percent of the genes showed increase in variability with age. However, we noted that the changes in variability calculated for each gene, using regression- and SVA-corrected data, were only weakly correlated, ρ(Δvar(ρ_regres_), Δvar(ρ_SVA_)) = 0.35 (Figure 3C).

**Figure 3.**
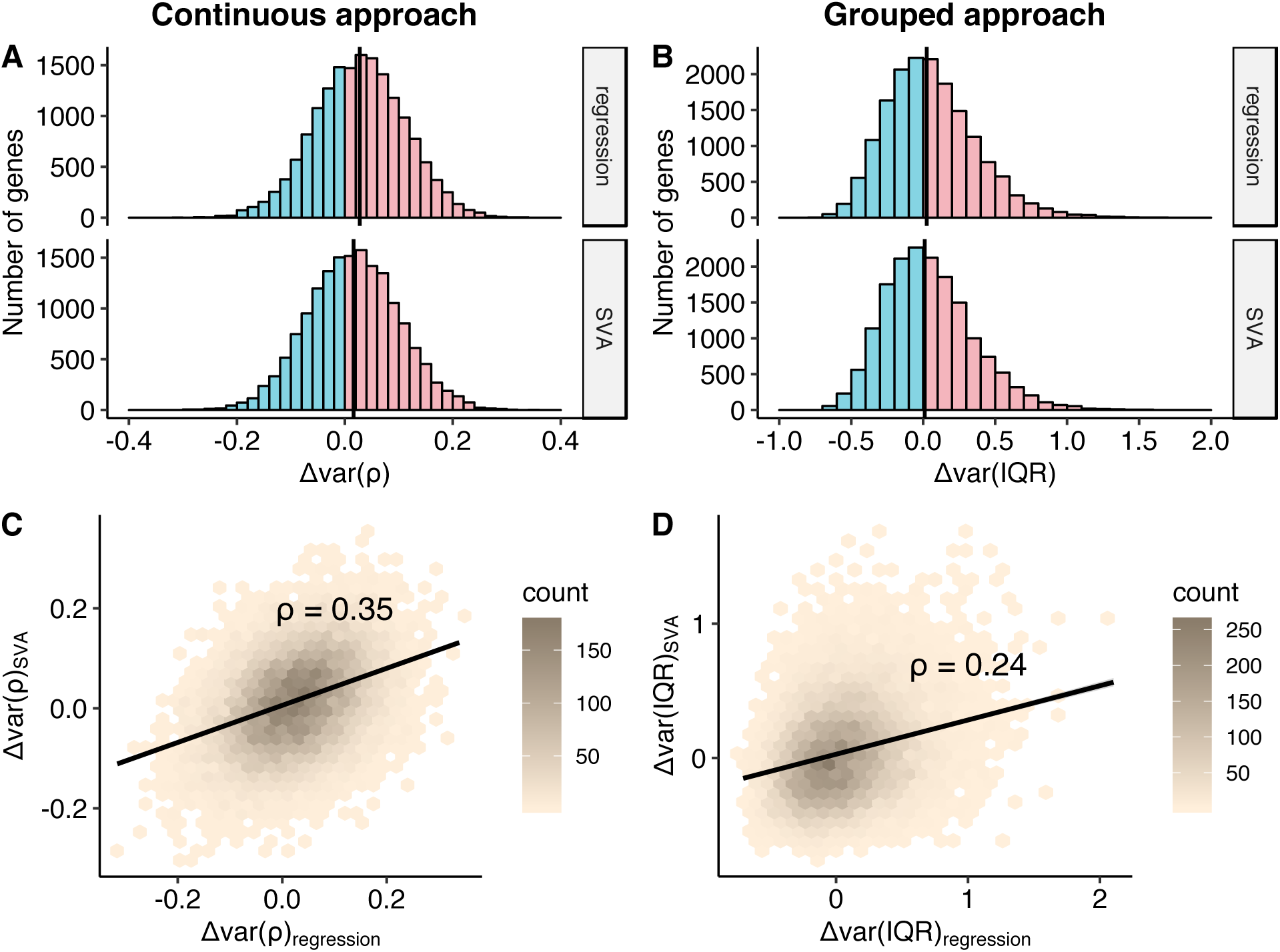
Comparison of the change in the variability addressed using continuous and grouped approaches, regression and SVA correction. Distributions of the Δvar-measures for all the genes (n = 16675) obtained in the continuous (A) and grouped (B) approaches. Increase in the variability with age, Δvar > 0, is coloured in pink, while decrease in variability, Δvar < 0, is marked in blue. The black straight line depicts median of the distribution. The Δvar(ρ) distributions are normal with their mean and median values equal to 0.03 and 0.02 for regression and SVA, respectively; The Δvar(IQR) distributions are moderately skewed: skewness values are 0.66 and 0.68 for regression and SVA, respectively. The mean and median values of the Δvar(IQR) distribution are 0.05 and 0.02 for regression and 0.04 and 0.01 for SVA, respectively. Hexagonal heat maps illustrate relationship between regression and SVA-corrected measures of the variability for each gene, obtained in continuous - Δvar(ρ) (C) and grouped - Δvar(IQR) (D) approaches. The colour gradient represents the density of the data. The linear regression line and the Spearman correlation estimate, ρ, for the corresponding variables are shown on each graph.

*In the grouped approach*, we first generated a distribution of expected variability in gene expression for the young individuals and treated it as a null distribution to compare with the variability from the old individuals. We used interquartile range (IQR) as a measure of variability, because it is robust to outliers. In order to calculate a distribution of expected variability in the young group, we randomly selected a subsample of 22 individuals (the number of samples in old group) from the 53 individuals in the young group 10 000 times and calculated IQR. The change in variability, Δvar (IQR), was measured as a fractional change in the IQR between old and young groups (see Methods). The p-value was determined by calculating how many times we observed a value as extreme as IQR_old_ (see Methods). The distributions of change in variability, Δvar(IQR), were moderately skewed to the right and ranged from −0.70 up to 2.10 for the regression corrected data and from −0.78 up to 1.71 for the SVA corrected data (Figure 3B). The skew to the right was expected given that we calculate variability change as a fraction and, thus, it was more sensitive to increase in variability. In both cases the distributions demonstrated a significant deviation from zero (Wilcox test, p value < 2.2e-16 both for regression and SVA corrections). The data revealed that 6% and 2% more genes showed more variability in the old group, for regression and SVA approaches respectively. Similar to the continuous approach, the effect sizes calculated using regression and SVA corrected data correlated weakly ρ(Δvar(IQR_regres_), Δvar(IQR_SVA_)) = 0.24, Figure 3D).

### Gene-level differential variability

We then asked if we could detect any genes with a significant change in variability. Using *the continuous approach*, we did not detect any significant change in variability with age after the multiple testing correction (Supplemental Table 3). *The grouped approach* leads to 741 and 746 differentially variable (DV) genes (FDR corrected p ≤ 0.05) using the regression and SVA correction, respectively (Figure 4A, Supplemental Table 4). However, the two sets of DV genes identified only have 83 genes in common (Figure 4A), one of which shows an opposite direction of change in the two sets. The correlation between Δvar (IQR) for regression and SVA corrected data is weak (ρ = 0.24), but correlation increases when we select only the common DV genes (ρ = 0.44) (Figure 4B). In agreement with our overview analysis above, we find twice as many DV genes with an increase in variability as those that decrease variability, using both correction methods: i) 533 genes increase and 208 decrease their variability in the regression correction, ii) 505 genes increase and 241 decrease their variability in the SVA correction (Figure 4A).

**Figure 4.**
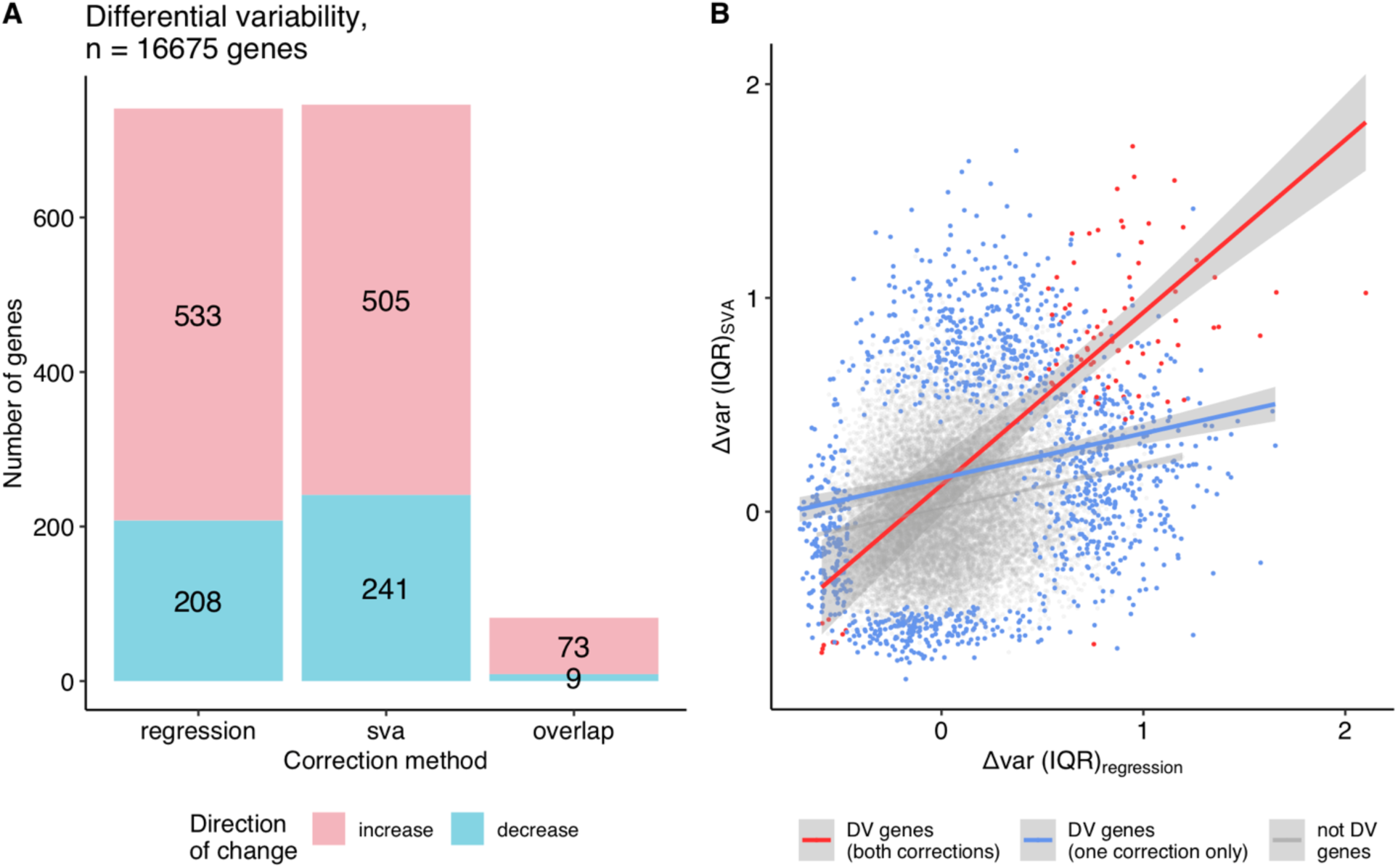
Differentially variable genes (grouped approach). (**A)** A bar plot of the differentially variable genes in ageing identified using the grouped approach (regression, SVA correction and their overlap), direction of the change in the variability is shown in colour: red - increase in variability in ageing, blue – decrease in variability (The single common gene, which shows inconsistency between approaches, is not displayed on the graph). (**B)** The relationship between the variability measures calculated using the grouped approach, Δvar (IQR), for regression and SVA-corrected data. Differentially variable (DV) genes identified in both corrections are highlighted in red (Spearman ρ = 0.44); DV genes identified in either regression, or SVA - in blue (Spearman ρ = 0.20); genes, that were not found to be differentially variable in any of the approaches – in grey (Spearman ρ = 0.20).

### Differential variability of functional groups

Following the individual gene analysis, we explored whether genes that tend to increase or decrease variability with age are localized in particular functional groups. We performed multiple gene set enrichment analyses (GSEA) using the change in the variability with age (Δvar) measures obtained in the continuous and grouped approaches on the gene sets from KEGG and Biological Process GO categories (Supplemental Tables 5-6). We observed no genome-level significant enrichment in particular functional groups on the data either from the continuous (SVA correction), or the grouped approach (Regression and SVA corrections). However, we found that 4 pathways, namely beta-Alanine metabolism, Ras signalling pathway, Phosphatidylinositol signalling system, Bacterial invasion of epithelial cells (FDR corrected p ≤ 0.05) were enriched among the genes showing more variability of expression in the continuous approach (Regression correction). These pathways had positive normalized enrichment scores (NES) i.e. enrichment for the genes that increase variability with age. Moreover, these pathways also had positive NES for other approaches, even though they were not significant (Supplemental Table 5).

### Distribution of the DV genes in the pathways

The gene set enrichment analysis shows if there are particular gene sets that include the genes with the highest increase or decrease. Failing to detect such functional categories, we asked how the variability measures for the genes were distributed in the different functional groups of genes. For each of 310 KEGG pathways, encompassing 5922 unique genes, we analysed the distributions of Δvar measures (Supplemental Table 7), focusing on the median value for the change in variability (Figure 5A, B). In line with the overall tendencies we observed (Figure 3A, B), the majority of pathways contained a larger number of genes that become more variable with age, irrespective of the approach or correction method used. Although the increase in variability is ubiquitous and is observed across the majority of the pathways (74-94%), the increase is small – in accordance with the small, but significant increase observed in the distribution for all genes. Since the pathways are not mutually exclusive, we checked if there are particular genes that are present in many different pathways and cause the shift. However, no significant correlation between the pathway membership of gene and its variability measure (Δvar) was detected (Supplemental Figure 11). We repeated the analysis using GO Biological Process categories and observed a similar trend (see Supplemental Information, Supplemental Table 7).

**Figure 5.**
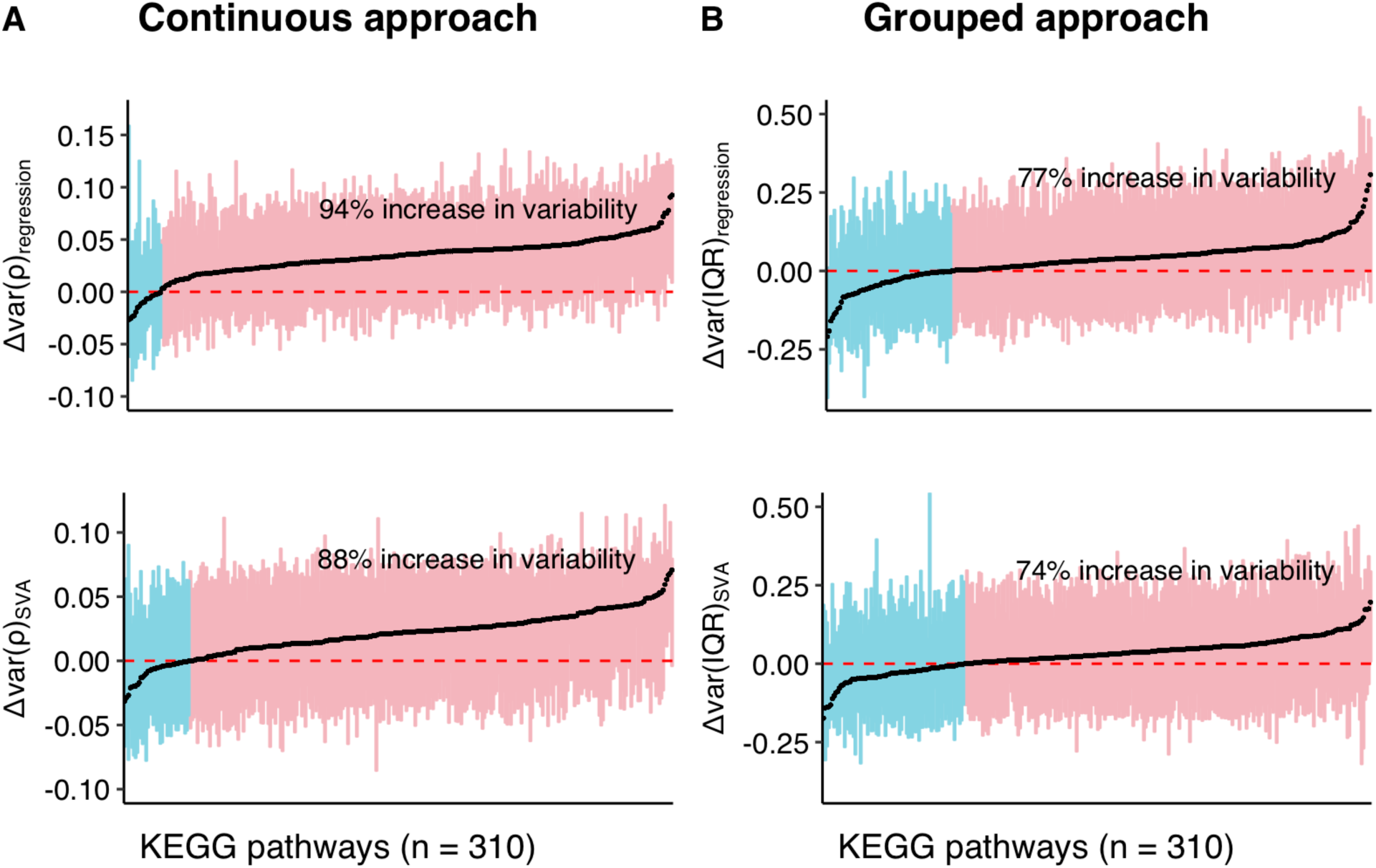
Distributions of the variability measures (Δvar) obtained using a combination of continuous (A) and grouped (B) approaches with regression and SVA-correction for the individual pathways in KEGG database. The distribution of the variability measures (Δvar) for the genes within each pathway is represented as a box, encapsulating part of the distribution between 1st and 3rd quantile, median of the box is coloured in black. Pathways on Y-axis and corresponding them boxes are ordered by increasing median. Boxes are coloured in red, if the corresponding pathways have median Δvar > 0, and in blue, if median Δvar < 0. Text label on the plot shows percentage of pathways with median Δvar > 0. Red dashed line marks Δvar = 0, while black straight-line marks median across all the pathways.

## DISCUSSION

Using one of the largest publicly available human brain expression datasets, we have investigated the change in the variability of the gene expression with age. We applied and compared different approaches to identify differentially variable genes and correction strategies to adjust for the confounders. Our comparison showed that the correction strategy plays a pivotal role in identifying the specific set of differentially variable (DV) genes. However, irrespective of the approach and correction method used, we observed a transcriptome-wide increase in the gene expression variability, i.e. more genes showed a tendency to increase than to decrease expression variability with age. We also showed that most of the functional processes (as defined in KEGG and GO) were susceptible to the ageing-related increase in the expression variability.

The difference between the *continuous* and *grouped approaches* can be explained by the power and initial assumptions of each method. While the *continuous approach* assumes a linear change in expression with age^0.2^ (see Supplemental Figure 4 for the results showing high concordance between models using age vs. age^0.2^ – correlation coefficient (ρ) ranges between 0.995 to 0.997), the *grouped approach* analyses each age-group within itself and is not sensitive to different dynamics of gene expression change. However, the *grouped approach* requires expression levels to be similar within the young and old groups. The *continuous approach* is well suited to detect monotonic changes in variability, whereas the *grouped approach* can detect switch-like changes, e.g. when variability stays the same throughout the lifespan but changes abruptly at the age of 60. In contrast, the *continuous approach* focuses on the whole ageing period, while the *grouped approach* overlooks the middle-age group (40-60). Finally, both methods are vulnerable to power issues as the *continuous approach* uses Spearman correlation, a non-parametric method, and the *grouped approach* analyses only a subset of the data. Thus, we compared the variability measure of each gene, calculated using these two approaches. The variability measures are moderately correlated (ρ = 0.43) for the regression correction and strongly correlated (ρ = 0.71) for the SVA correction (Supplemental Figure 9). Overall, the differences in the results using these two approaches create a challenge in interpretation, but they are not surprising given inherent differences in methodology and the small changes in variability we are investigating.

Another technical aspect we considered was the effect of pre-processing steps. While applying regression and SVA corrections, we showed that significantly DV genes hardly overlap between the corrections, with only 6% being in common (Jaccard similarity) (Figure 4A). Unfortunately, current approaches for handling transcriptome data are designed only to remove the confounding factors on the expression level and not on the expression variability. Thus, SVA and regression demonstrated much higher agreement in the differentially expressed (DE) genes (24% in common, Jaccard similarity) (Figure 1B). That raises a question: which set includes the genuine DV genes? The different correction strategies are quite distinct and might be accounting for different aspects, which is evident from the weak correlation between them (Spearman ρ between regression and SVA-corrected data for continuous approach – 0.35, grouped approach − 0.24, Supplemental Figure 6-7). In this case, the union may capture the full aspects of differential variability, whereas the overlap can provide the gene list in which we are most confident.

Independent of the correction strategy, two thirds of the DV genes showed a significant increase in variability. These results agree with the reports of Li *et al.* [10] on mice neural retina, but disagree with findings of Brinkmeyer-Langford *et al.* [11] on the human brain and Vinuela *et al.* [17] on the multiple human tissues which show either equal amount of genes increase and decrease in variability or more decrease than increase. The small overlap of our DV gene set with Brinkmeyer-Langford *et al.* [11] (see Supplemental Information) could be explained by the technical aspects that we presented, i.e. variability measure and data pre-processing, as well as use of different experimental setups and different age-ranges.

We further asked if there is a shift towards an increase or decrease in variability (above or below zero) across the whole transcriptome, irrespective of the values and significance. In accordance with the previous findings on the human, rat and fruit fly [6,12,15], we found as many as 63% of genes showed increase in variability, whereas the value was lower for the grouped approach, i.e. 51%. Functional investigation of the differential variability showed that it is ubiquitous and was not concentrated in specific functional groups. That was further supported by the fact that as many as 74% to 94% of KEGG pathways included more genes with an increase in variability (Figure 5 A, B).

Most studies consider the accumulation of cellular damage, such as somatic mutations, with age as a main factor, causing increase in the gene expression variability with age. Indeed, Lodato *et al* [26] shows increase in the number of single nucleotide variants in human brain with age, while Lee *et al* [27] documented somatic recombination of *APP* gene in human neurons and its increase with age. However, the causal link between the accumulation of mutations and increase in variability was not proven and Enge *et al.* [15] provides an evidence that somatic mutations are not enough to explain gene expression variability. Moreover, because brain is a post-mitotic tissue, it may demonstrate a different damage profile, as it is not as prone to replication-associated mutations as other tissues but associated with other types of damage, such as free radicals or loss of proteostasis. Since we analyse expression values from different individuals, we should consider the effect of genotype. A few studies have identified a small set of genetic variants that could change gene expression during ageing (genotype-by-age interaction) [11,17,28]. However, these specific differences in genotype are not likely on their own to explain the transcriptome-wide shift that we observed. The environmental factors influencing the epigenome, as well as stochastic effects driving an epigenetic drift [9,17,18,29] seems to be a likely explanation in this case. The change in variability could also stem from the change in gene expression levels. Although not replicated in mouse brain [14], Davie *et al.*[12] shows that ageing leads to an overall decrease in the RNA content, which could also be the reason for such a global increase in the expression variability. However, we apply log2 transformation, which attempts to correct the mean-variability dependence. Indeed, we do not observe any significant association between the changes in expression level and variability (Odds ratio = 1.05, Fisher’s test p = 0.11, Supplemental Figure 10).

Although we used one of the largest, well-characterized datasets, it is important to note that the sample size, the unequal coverage of ages and the high technical and biological variation all posed a challenge for the analysis. Moreover, this data was generated using microarray technology, which does not measure the expression of all genes and is not as quantitative as RNA-seq. Future studies addressing variability in gene expression may consider use of scRNA-seq data to distinguish unique changes within a cell from the coordinated changes within cell population or changes in the cell composition.

Providing a systematic analysis of the same dataset at multiple levels and considering multiple technical challenges, we showed a slight but significant shift towards an age-related increase in variability that was not clustered in certain functions but distributed across all pathways. It has been recently suggested that an increase in expression variability is linked with the genetic risk for schizophrenia in males [30]. However, future experiments are crucial to understand whether all genes, functions and organs are equally tolerant to the variability we observed and whether this variability has any causal relationship with the ageing processes.

## METHODS

### Data processing steps

#### Dataset Selection

We utilized one of the largest age-series human brain expression datasets, featuring 269 prefrontal cortex samples from healthy individuals and spanning the whole lifespan from development (prenatal samples) through ageing (80 years) [23]. These data were collected using microarray technology from people of both sexes and 4 races, namely African American (AA), Caucasian (CAUC), Hispanic (HISP) and Asian (AS). In the current analysis we excluded foetal, childhood and early adulthood samples before the age of 20, thus limiting our sample size to 147. This was to exclude developmental processes taking place in the brain until the end of early adulthood, which exhibit discontinuous expression changes between early adulthood and ageing [31]. Our main motivation was to study changes in gene expression variability during ageing, considering 20 years old as a starting point.

#### Data Characterization

The pre-processed data (loess normalization was applied on the background corrected log2 intensity ratios (sample/reference)[23]); sample and gene (probe set to Entrez gene mapping) annotations were obtained from the NCBI Gene Expression Omnibus (GEO) at accession number GSE30272. Samples were processed in 19 batches, had different quality measurements, namely pH and RNA integrity number (RIN), and differed in the time of collection after death (post-mortem interval (PMI)). Using a PCA, we found no sample outliers as judged by visual inspection of the first two principal components (Supplemental Figure 2). However, the relationship analysis between the above-mentioned factors (i.e. batch, RIN, PMI and others) and age yielded significant correlations for sex, post-mortem interval and RNA integrity, pointing to potential confounders in the data (Supplemental Figure 1). We further checked the overlap between significantly differentially variable genes in our analysis and previously reported genes that are affected by PMI and detected only a limited overlap (see Supplemental Information).

#### Probe set to Gene summarization

If one probe-set was mapped to several genes, it was deleted to avoid duplication. Conversely, when one gene had several probe-set expression values, they were averaged to obtain a unique gene expression value. In total 16675 genes were measured on the array.

#### Batch correction

To compensate for technical variation between samples, quantile normalisation (QN) was performed using the ‘normalise.quantiles’ function from the ‘preprocessCore’ R library. To differentiate between the age effect and the effect of the unwanted technical and biological variability, we have applied different expression correction strategies: linear regression of the known covariates, unsupervised estimation of covariates using surrogate variable analysis (SVA) [25,32], and ComBat, a parametric empirical Bayesian framework for covariate adjustment [24]. As a result, we analyzed the same data four times, corrected using QN, QN+regression, QN+SVA, QN+ComBat. Different corrections work by adjusting for the different covariates in the linear model that explains the gene expression, namely: i) QN – no covariates were added; ii) QN+regression – 25 covariates considered: technical batches (N = 19), sex (N=2), race (N=4), post-mortem interval, RNA integrity number,pH; iii) QN + SVA – 20 surrogate variables (SV) were inferred from the expression data using the ‘sva’ function from “SVA” R library; iv) QN+ComBat: the 6 confounding factors: batch (N = 19), sex (N=2), race(N=4), pmi, RIN and pH were adjusted for, one at the time, by repeatedly applying the ComBat function from the “SVA” R library to the expression data.

### Differential expression

A least squares linear regression model was used to model gene expression level change with age. Age^0.25^ was used as an independent variable instead of age to account for the difference in rate of gene expression changes between young (fast) and old (slow) as well as different density of the samples across ages. Nevertheless, the β_1_-coefficients from the linear model, that uses age^0.25^ correlate well with the one, that employs age (Supplemental Figure 4). Coefficients for the age covariate were used as a measure of the differential expression. P values for coefficients were adjusted using the FDR method with a threshold p ≤ 0.05 to account for multiple testing. Depending on the correction method applied, the linear model also accounted for different measured or unmeasured covariates (see Data processing steps) of the following general form:

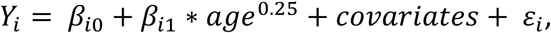

where *Y*_i_ is the normalized log-expression level of a gene with i = 1,…,n, β_i0_-intercept, β_i1_-slope term and ε_i_ - residual (or error) term.

### Differential variability

#### The continuous approach

First, a linear model to fit gene expression during ageing, using age^0.2^ and potential confounders, was constructed. Next, the Spearman correlation was calculated between the absolute values of the residuals, |ε_i_| from the linear model and age. Consequently, Spearman correlation estimates were used as a measure of the change in variability, referred as Δvar_i_(ρ). P values for the Spearman correlation estimates were corrected for multiple testing using FDR. FDR adjusted p ≤ 0.05 was used as a threshold to define significantly DV genes.

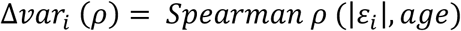

#### The grouped approach

First, a corrected expression matrix was obtained by removing the effect of covariates (see data processing steps) from the data using the residuals from a linear regression model (Y_i_ = β_i0_ + covariates + ε). The ‘grouped approach’ is a custom resampling-based test designed to compare gene expression variability between young (20 – 40 years old) and old (60-80 years old) groups using an interquartile range (IQR). IQR corresponds to the difference between the 75^th^ and 25^th^ percentiles of the distribution and is considered to be a robust measure of variability, meaning it is not susceptible to outliers and departure from normality in the data. In order to adjust for the unequal sample size of the young (N = 53) and old (N = 22) groups, we, first, calculated a null distribution of the IQR values for the young group by resampling it 10 000 times with the size of the old group. Next, we calculated significance as a percentage of samples where IQR_old_ was more extreme than IQR_young_ and corrected it for multiple testing using FDR correction, q ≤ 0.05. The ‘grouped’ measure of change in the variability, Δvar_i_(IQR), for the gene *i*, corresponds to the difference between IQR value for the old,*IQR_i old_* and 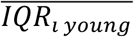 (i.e. mean IQR value from the young distribution), which is then divided by the latter, see formula:

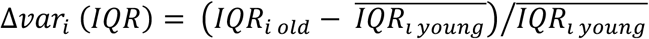

### Gene Set Enrichment Analysis for KEGG pathways and GO categories

β_1_ coefficients from the differential expression and Δvar measures from the differential variability analyses were used to perform gene set enrichment analysis, GSEA [33] using the “clusterProfiler” R library. KEGG pathways (N = 315) and BP GO terms (Biological processes Gene Ontology, N = 5822) with the size of between 10 and 500 genes were considered as gene sets for the GSEA.

### Pathway Distribution Study

KEGG pathway to gene mapping was obtained from “KEGGREST” R library and pathways were pre-filtered to contain between 5 and 500 genes. As a result, 310 KEGG pathways that comprise 5922 unique genes were used for the subsequent analysis. The boxplots illustrated distributions of the Δvar measure for genes in each pathway. Pathways were sorted according to their median Δvar measure in ascending order. The percentage of pathways that have their median Δvar above zero was calculated. The analysis was replicated using BP GO terms (N = 5919) of a size between 10 and 500 genes, which in total contained 12538 unique genes. Mapping of GO terms to genes was obtained from “org.Hs.eg.db” R library.

### Distributions tests

Distributions of the Δvar - measures for all the genes were tested for normality using the Shapiro-Wilk test in R (‘Shapiro.test’ function) on the multiple subsamples, consisting of 5000 measures. Skewness of the distributions was calculated using the ‘fBasics’ function from “BasicStatistics” R library.

### Mean-variability relationship testing

To visualise and test if the change in gene expression variability is associated with the change in gene expression level, we plotted the difference in the means between the young and old groups against difference in the interquartile range (IQR) between the young and old groups. Mean and IQR for the old group were calculated once, while mean and IQR for the young group were calculated 10,000 times for the subsamples (see Grouped approach) and then means of the distributions of the corresponding values (mean and IQR) were used in the analysis. Fisher’s exact test was performed on the values used for the plotting.

### Software

R version 3.5.0 and “data.table” were used to perform the analyses, while “ggplot2” and “ggpubr” R libraries were used to create visualizations of the data.

## ABBREVIATIONS

BP GO: Biological Process Gene Ontology
DE: differentially expressed genes
DV: differentially variable genes
Δvar: measure of change in the expression variability with age
GO: Gene Ontology
GSEA: Gene Set Enrichment Analysis
IQR: interquartile range
KEGG: Kyoto Encyclopedia of Genes and Genomes
PMI: post-mortem interval
QN: quantile normalization
RIN: RNA integrity number
Rho: Spearman correlation estimate
SVA: Surrogate Variable Analysis

## AUTHOR CONTRIBUTIONS

H.M.D, V.R.K and J.M.T designed the study. V.R.K and H.M.D analysed the data. V.R.K, H.M.D and J.M.T interpreted the results and wrote the manuscript. All authors read, revised and approved the final version of this manuscript.

## ACKNOWLEDGEMENTS

The authors thank to Dr. Mehmet Somel, Dr. John Marioni and the members of Thornton group, especially Dr. Dobril Ivanov and Daniel Elias Martin Herranz for their support and helpful discussions and to Matías Fuentealba Valenzuela for critical reading of the manuscript. H.M.D is a member of Darwin College, University of Cambridge.

## REFERENCES

1. Herndon LA, Schmeissner PJ, Dudaronek JM, Brown PA, Listner KM, Sakano Y, Paupard MC, Hall DH, Driscoll M. Stochastic and genetic factors influence tissue-specific decline in ageing C. elegans. Nature [Internet]. 2002; 419: 808–14. Available from: http://dx.doi.org/10.1038/nature01135

2. Kirkwood TBL, Feder M, Finch CE, Franceschi C, Globerson A, Klingenberg CP, LaMarco K, Omholt S, Westendorp RG. What accounts for the wide variation in life span of genetically identical organisms reared in a constant environment? Mech Ageing Dev [Internet]. Elsevier; 2005; 126: 439–43. Available from: https://doi.org/10.1016/j.mad.2004.09.008

3. Horvath S. DNA methylation age of human tissues and cell types. Genome Biol [Internet]. 2013; 14: R115. Available from: https://doi.org/10.1186/gb-2013-14-10-r115

4. Fraga MF, Ballestar E, Paz MF, Ropero S, Setien F, Ballestar ML, Heine-Suner D, Cigudosa JC, Urioste M, Benitez J, Boix-Chornet M, Sanchez-Aguilera A, Ling C, et al. Epigenetic differences arise during the lifetime of monozygotic twins. Proc Natl Acad Sci [Internet]. 2005; 102: 10604–9. Available from: https://doi.org/10.1073/pnas.0500398102

5. Cheung P, Vallania F, Warsinske HC, Donato M, Schaffert S, Chang SE, Dvorak M, Dekker CL, Davis MM, Utz PJ, Khatri P, Kuo AJ. Single-Cell Chromatin Modification Profiling Reveals Increased Epigenetic Variations with Aging. Cell [Internet]. Elsevier; 2018 [cited 2018 Sep 21]; 173: 1385–1397.e14. Available from: https://doi.org/10.1016/j.cell.2018.03.079

6. Somel M, Khaitovich P, Bahn S, Pääbo S, Lachmann M. Gene expression becomes heterogeneous with age. Curr Biol [Internet]. Elsevier; 2006; 16: R359–60. Available from: https://doi.org/10.1016/j.cub.2006.04.024

7. Bahar R, Hartmann CH, Rodriguez KA, Denny AD, Busuttil RA, Dollé MET, Calder RB, Chisholm GB, Pollock BH, Klein CA, Vijg J. Increased cell-to-cell variation in gene expression in ageing mouse heart. Nature [Internet]. 2006; 441: 1011–4. Available from: http://dx.doi.org/10.1038/nature04844

8. Martinez-Jimenez CP, Eling N, Chen H-C, Vallejos CA, Kolodziejczyk AA, Connor F, Stojic L, Rayner TF, Stubbington MJT, Teichmann SA, de la Roche M, Marioni JC, Odom DT. Aging increases cell-to-cell transcriptional variability upon immune stimulation. Science (80-) [Internet]. 2017; 355: 1433–6. Available from: https://doi.org/10.1126/science.aah4115

9. Hernando-Herraez I, Evano B, Stubbs T, Commere P-H, Clark S, Andrews S, Tajbakhsh S, Reik W. Ageing affects DNA methylation drift and transcriptional cell-to-cell variability in muscle stem cells. bioRxiv [Internet]. Cold Spring Harbor Laboratory; 2018 [cited 2019 Jan 16];: 500900. Available from: https://doi.org/10.1101/500900

10. Li Z, Wright FA, Royland J. Age-dependent variability in gene expression in male fischer 344 rat retina. Toxicol Sci [Internet]. Oxford University Press; 2009 [cited 2018 May 1]; 107: 281–92. Available from: https://doi.org/10.1093/toxsci/kfn215

11. Brinkmeyer-Langford CL, Guan J, Ji G, Cai JJ. Aging Shapes the Population-Mean and - Dispersion of Gene Expression in Human Brains. Front Aging Neurosci [Internet]. Frontiers; 2016 [cited 2019 Jan 21]; 8: 183. Available from: https://doi.org/10.3389/fnagi.2016.00183

12. Davie K, Janssens J, Koldere D, De Waegeneer M, Pech U, Kreft Ł, Aibar S, Makhzami S, Christiaens V, Bravo González-Blas C, Poovathingal S, Hulselmans G, Spanier KI, et al. A Single-Cell Transcriptome Atlas of the Aging Drosophila Brain. Cell [Internet]. 2018; 174: 982–998.e20. Available from: https://doi.org/10.1016/j.cell.2018.05.057

13. Angelidis I, Simon LM, Fernandez IE, Strunz M, Mayr CH, Greiffo FR, Tsitsiridis G, Graf E, Strom TM, Eickelberg O, Mann M, Theis FJ, Schiller HB. An atlas of the aging lung mapped by single cell transcriptomics and deep tissue proteomics. bioRxiv [Internet]. Cold Spring Harbor Laboratory; 2018 [cited 2019 Jan 16];: 351353. Available from: https://doi.org/10.1101/351353

14. Ximerakis M, Lipnick SL, Simmons SK, Adiconis X, Innes BT, Dionne D, Nguyen L, Mayweather BA, Ozek C, Niziolek Z, Butty VL, Isserlin R, Buchanan SM, et al. Single-cell transcriptomics of the aged mouse brain reveals convergent, divergent and unique aging signatures. bioRxiv [Internet]. Cold Spring Harbor Laboratory; 2018 [cited 2019 Jan 16];: 440032. Available from: https://doi.org/10.1101/440032

15. Enge M, Arda HE, Mignardi M, Beausang J, Bottino R, Kim SK, Quake SR. Single-Cell Analysis of Human Pancreas Reveals Transcriptional Signatures of Aging and Somatic Mutation Patterns. Cell [Internet]. Elsevier; 2017 [cited 2017 Oct 3]; 171: 321– 330.e14. Available from: https://doi.org/10.1016/j.cell.2017.09.004

16. Wiley CD, Flynn JM, Morrissey C, Lebofsky R, Shuga J, Dong X, Unger MA, Vijg J, Melov S, Campisi J. Analysis of individual cells identifies cell-to-cell variability following induction of cellular senescence. Aging Cell [Internet]. 2017 [cited 2017 Jul 14]; 16: 1043–50. Available from: http://doi.wiley.com/10.1111/acel.12632

17. Viñuela A, Brown AA, Buil A, Tsai P-C, Davies MN, Bell JT, Dermitzakis ET, Spector TD, Small KS. Age-dependent changes in mean and variance of gene expression across tissues in a twin cohort. Hum Mol Genet [Internet]. Cold Spring Harbor Laboratory; 2018 [cited 2017 Sep 21]; 27: 732–41. Available from: https://doi.org/10.1093/hmg/ddx424

18. Slieker RC, van Iterson M, Luijk R, Beekman M, Zhernakova D V., Moed MH, Mei H, van Galen M, Deelen P, Bonder MJ, Zhernakova A, Uitterlinden AG, Tigchelaar EF, et al. Age-related accrual of methylomic variability is linked to fundamental ageing mechanisms. Genome Biol [Internet]. BioMed Central; 2016 [cited 2017 Sep 21]; 17: 191. Available from: https://doi.org/10.1186/s13059-016-1053-6

19. Inukai S, Pincus Z, de Lencastre A, Slack FJ. A microRNA feedback loop regulates global microRNA abundance during aging. RNA [Internet]. Cold Spring Harbor Laboratory Press; 2018 [cited 2018 May 1]; 24: 159–72. Available from: https://doi.org/10.1261/rna.062190.117

20. Warren LA, Rossi DJ, Schiebinger GR, Weissman IL, Kim SK, Quake SR. Transcriptional instability is not a universal attribute of aging. Aging Cell [Internet]. 2007; 6: 775–82. Available from: https://doi.org/10.1111/j.1474-9726.2007.00337.x

21. Southworth LK, Owen AB, Kim SK. Aging Mice Show a Decreasing Correlation of Gene Expression within Genetic Modules. PLoS Genet [Internet]. Public Library of Science; 2009; 5: e1000776. Available from: https://doi.org/10.1371/journal.pgen.1000776

22. Oh G, Ebrahimi S, Wang SC, Cortese R, Kaminsky ZA, Gottesman II, Burke JR, Plassman BL, Petronis A. Epigenetic assimilation in the aging human brain. Genome Biol [Internet]. BioMed Central; 2016 [cited 2018 May 1]; 17: 76. Available from: https://doi.org/10.1186/s13059-016-0946-8

23. Colantuoni C, Lipska BK, Ye T, Hyde TM, Tao R, Leek JT, Colantuoni EA, Elkahloun AG, Herman MM, Weinberger DR, Kleinman JE. Temporal dynamics and genetic control of transcription in the human prefrontal cortex. Nature [Internet]. Nature Publishing Group, a division of Macmillan Publishers Limited. All Rights Reserved.; 2011; 478: 519–23. Available from: http://dx.doi.org/10.1038/nature10524

24. Johnson WE, Li C, Rabinovic A. Adjusting batch effects in microarray expression data using empirical Bayes methods. Biostatistics [Internet]. Oxford University Press; 2007; 8: 118–27. Available from: https://doi.org/10.1093/biostatistics/kxj037

25. Leek JT, Storey JD. Capturing Heterogeneity in Gene Expression Studies by Surrogate Variable Analysis. PLoS Genet [Internet]. Chapman & Hall; 2007 [cited 2017 Jul 27]; 3: e161. Available from: https://doi.org/10.1371/journal.pgen.0030161

26. Lodato MA, Rodin RE, Bohrson CL, Coulter ME, Barton AR, Kwon M, Sherman MA, Vitzthum CM, Luquette LJ, Yandava CN, Yang P, Chittenden TW, Hatem NE, et al. Aging and neurodegeneration are associated with increased mutations in single human neurons. Science (80-) [Internet]. American Association for the Advancement of Science; 2018 [cited 2019 Jan 18]; 359: 555–9. Available from: https://doi.org/10.1126/science.aao4426

27. Lee M-H, Siddoway B, Kaeser GE, Segota I, Rivera R, Romanow WJ, Liu CS, Park C, Kennedy G, Long T, Chun J. Somatic APP gene recombination in Alzheimer’s disease and normal neurons. Nature [Internet]. Nature Publishing Group; 2018; 563: 639–45. Available from: https://doi.org/10.1038/s41586-018-0718-6

28. Kent JW, Göring HHH, Charlesworth JC, Drigalenko E, Diego VP, Curran JE, Johnson MP, Dyer TD, Cole SA, Jowett JBM, Mahaney MC, Comuzzie AG, Almasy L, et al. Genotype×age interaction in human transcriptional ageing. Mech Ageing Dev [Internet]. Elsevier; 2012 [cited 2018 May 1]; 133: 581–90. Available from: https://doi.org/10.1016/j.mad.2012.07.005

29. Pal S, Tyler JK. Epigenetics and aging. Sci Adv [Internet]. American Association for the Advancement of Science; 2016 [cited 2019 Jan 21]; 2: e1600584. Available from: https://doi.org/10.1126/sciadv.1600584

30. Chen J, Cao H, Meyer-Lindenberg A, Schwarz E. Male increase in brain gene expression variability is linked to genetic risk for schizophrenia. Transl Psychiatry [Internet]. 2018; 8: 140. Available from: https://doi.org/10.1038/s41398-018-0200-0

31. Dönertaş HM, İzgi H, Kamacıoğlu A, He Z, Khaitovich P, Somel M. Gene expression reversal toward pre-adult levels in the aging human brain and age-related loss of cellular identity. Sci Rep [Internet]. 2017; 7: 5894. Available from: https://doi.org/10.1038/s41598-017-05927-4

32. Leek JT, Johnson WE, Parker HS, Jaffe AE, Storey JD. The sva package for removing batch effects and other unwanted variation in high-throughput experiments. Bioinformatics [Internet]. Oxford University Press; 2012; 28: 882–3. Available from: https://doi.org/10.1093/bioinformatics/bts034

33. Subramanian A, Tamayo P, Mootha VK, Mukherjee S, Ebert BL, Gillette MA, Paulovich A, Pomeroy SL, Golub TR, Lander ES, Mesirov JP. Gene set enrichment analysis: A knowledge-based approach for interpreting genome-wide expression profiles. Proc Natl Acad Sci [Internet]. 2005; 102: 15545–50. Available from: https://doi.org/10.1073/pnas.0506580102

