## Supplemental Information for "The variability of expression of many genes and most functional pathways is observed to increase with age in brain transcriptome data"

### SUPPLEMENTAL DATA

#### Supplemental Tables

**Supplemental Table 1**
The results of the differential expression analysis performed on regression-corrected and SVA-corrected data

**Supplemental Table 2**
The results of the GSEA Analysis for BP GO categories performed on the differentially expressed genes obtained on regression-corrected and SVA-corrected data

**Supplemental Table 3**
The results of the differential variability analysis while applying “continuous approach” for the regression and SVA-corrected expression data.

**Supplemental Table 4**
The results of the differential variability analysis while applying “grouped approach” for the regression and SVA-corrected expression data.

**Supplemental Table 5 and 6**
The results of GSEA Analysis among the KEGG pathways and GO categories (separate analysis was performed for different groups) on the differential variability measures from the “continuous” and “grouped” approaches.

**Supplemental Table 7**
Mapping of genes and their variability measures to KEGG pathways and GO terms that were used to calculate median statistics across the KEGG pathways and GO terms.

**Supplemental Information**

**Post-mortem interval associated differentially variable genes**

Recent reports made on the GTEx data [1] suggest that extent of Post-mortem interval (PMI) – the time passed between death and sample collection, could be associated with the degradation of specific mRNA species and, consequently, change in both mean and variability of their gene expression. We have checked whether our set of differentially variable genes (DV) contains any of the reported 266 genes that are susceptible to the degradation dependent on PMI. We found 14, 14, 10 and 7 of those genes among the set of differentially variable genes identified in the grouped approach with quantile normalisation, regression, ComBat and SVA correction, respectively.

**Overlap of the DV genes with other publications**

The total set of differentially variable genes identified by Brinkmeyer-Langford [2] comprises 848 distinct genes found across 13 brain regions. 171 of those genes are specific to the cortex and frontal cortex. Our set of DV genes found in grouped approach after regression correction (N total = 741) has an overlap of 21 genes with Brinkmeyer-Langford and 19 genes for the SVA-corrected expression (N total = 746). The percentage of overlap doesn't increase, if we only consider cortex-specific DV genes from Brinkmeyer-Langford - it constitutes 4 and 3 genes for regression and SVA corrected sets, respectively.

**Distribution of the differential variability in the KEGG pathways vs GO categories**

**Table 1 Percentage of KEGG pathways and GO terms that have median expression variability above 0, according to the different approaches**

| | Continuous approach | | | | Grouped approach | | | |
|---|---|---|---|---|---|---|---|---|
| | Regression | SVA | QN | ComBat | Regression | SVA | QN | ComBat |
| KEGG pathways | 94% | 88% | 99% | 88% | 77% | 74% | 94% | 95% |
| GO terms | 92% | 84% | 98% | 79% | 66% | 61% | 88% | 88% |

**References**


1. Zhu Y, Wang L, Yin Y, Yang E. Systematic analysis of gene expression patterns associated with postmortem interval in human tissues. Sci Rep [Internet]. Nature Publishing Group; 2017 [cited 2017 Sep 8]; 7: 5435. Available from: http://www.ncbi.nlm.nih.gov/pubmed/28710439
2. Brinkmeyer-Langford CL, Guan J, Ji G, Cai JJ. Aging Shapes the Population-Mean and -Dispersion of Gene Expression in Human Brains. Front Aging Neurosci [Internet]. Frontiers Media SA; 2016 [cited 2017 Sep 8]; 8: 183. Available from: http://www.ncbi.nlm.nih.gov/pubmed/27536236

#### Supplemental Figures

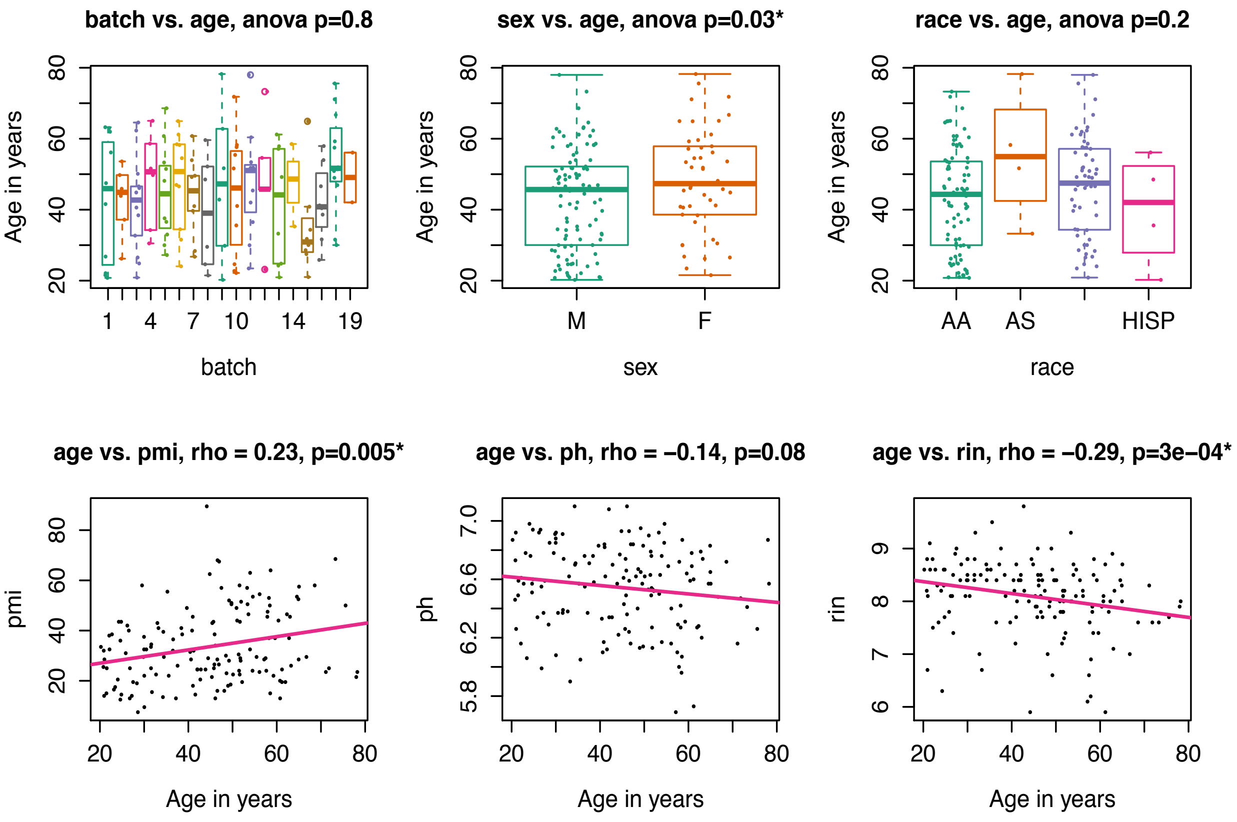


**Supplemental Figure 1.** Dependency plots between the age and different biological and technical factors, such as batch, sex, race, post-mortem interval (PMI), pH and RNA integrity number (RIN). Box plots were constructed for categorical covariates (batch, sex and race), while scatter plots were made for the numerical covariates (PMI, pH, RIN). In order to test for the significance of the dependence between categorical covariates and age ANOVA-test was performed (p value is displayed on the graph), Spearman correlation estimate was calculated to estimate relationship between numerical covariates and age (Spearman rho and p value are displayed on the graph). * $p \leq 0.05$

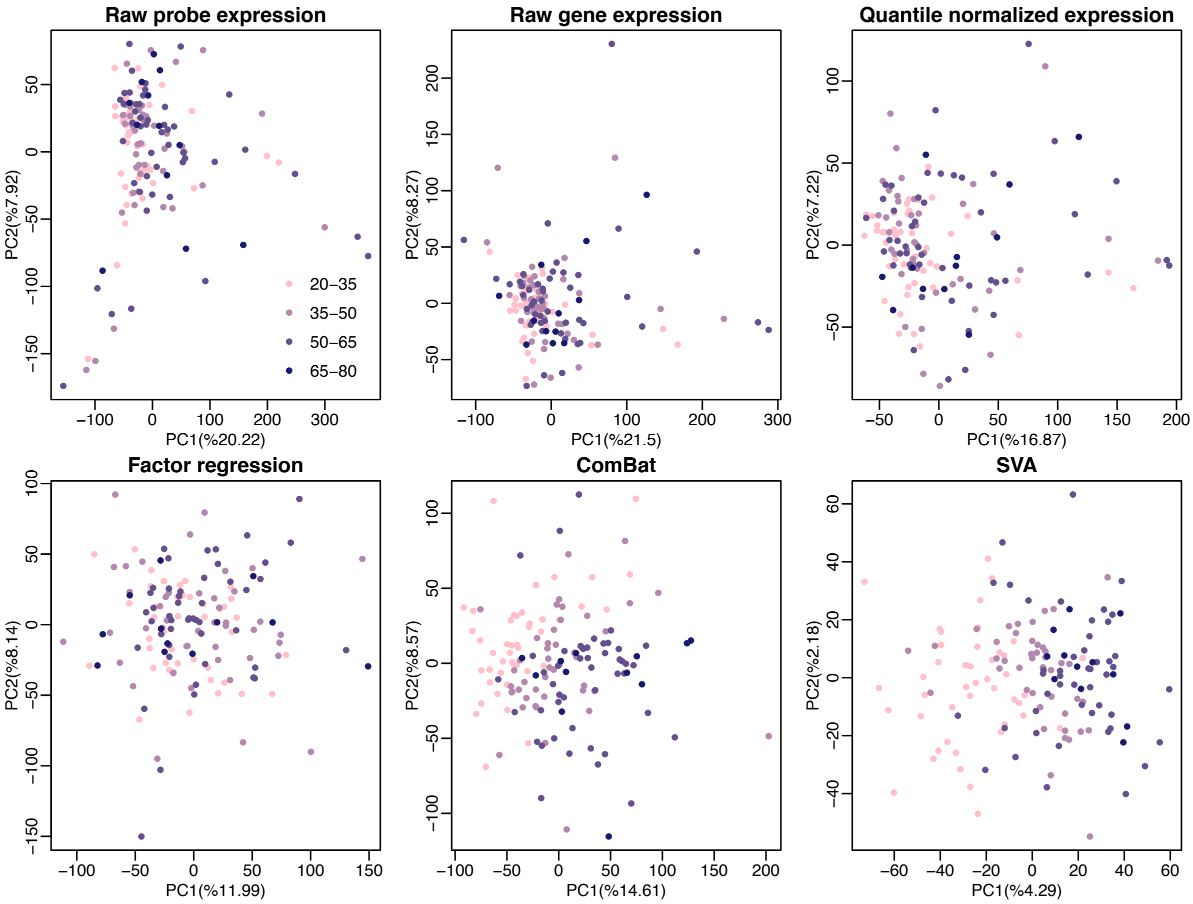


**Supplemental Figure 2.** PCA plots of the raw expression data and after different pre-processing steps. First row from left to right: probe expression level, gene expression level and quantile normalized gene expression level. Second row from left to right: quantile normalized data further corrected either with regression, ComBat or SVA. Colour gradient depicts age range from pink – young to violet – old.

**A**

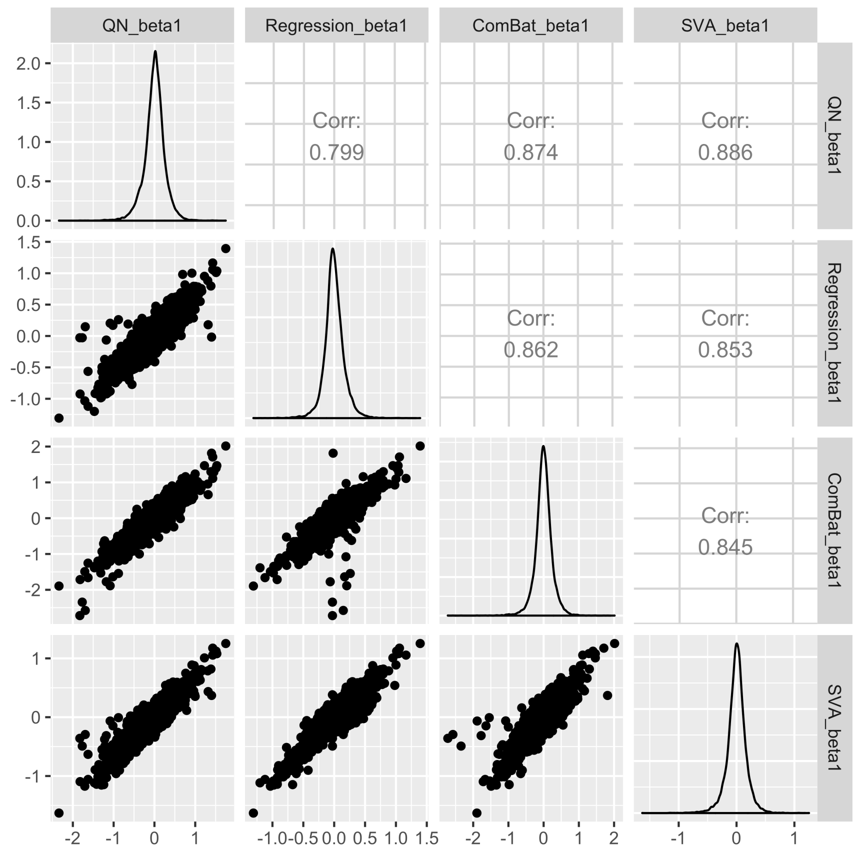


**B**

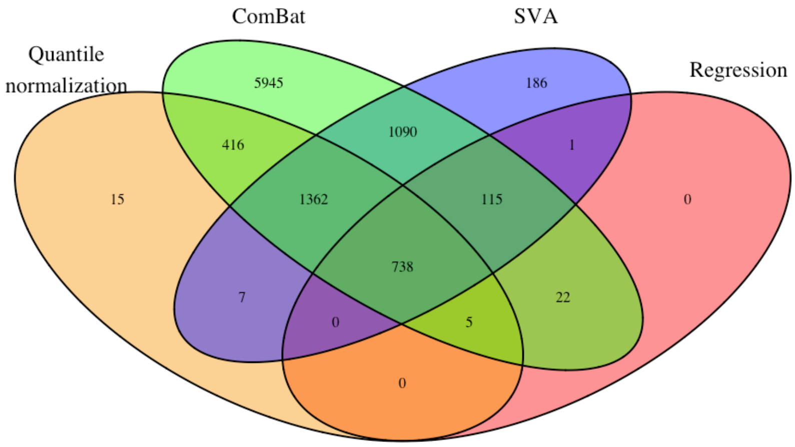


**Supplemental Figure 3. (A)** Illustration of the relationship between beta1 – coefficients from the linear regression models that explain gene expression corrected with different approaches (QN only, QN followed by regression, ComBat, SVA) using age$^{0.25}$. Scatter plots that illustrate relationship between the beta1 – coefficients from different corrections are displayed in the lower triangle of the matrix, distributions of the beta1 – coefficients are located on the diagonal and Spearman correlation estimates are displayed in the upper triangle of the matrix. **(B)** Venn diagram of the differentially expressed (DE) genes with age that were identified after applying different expression correction approaches (QN only, QN followed by regression, ComBat, SVA). The ComBat approach seem to greatly outnumber all the other approaches in the number of DE genes. We consider that consecutive application

(6 times) of the linear model that contains age and particular confounding factor caused an overfitting of gene expression with age and lead to identification of so many DE genes, many of which are false positives.

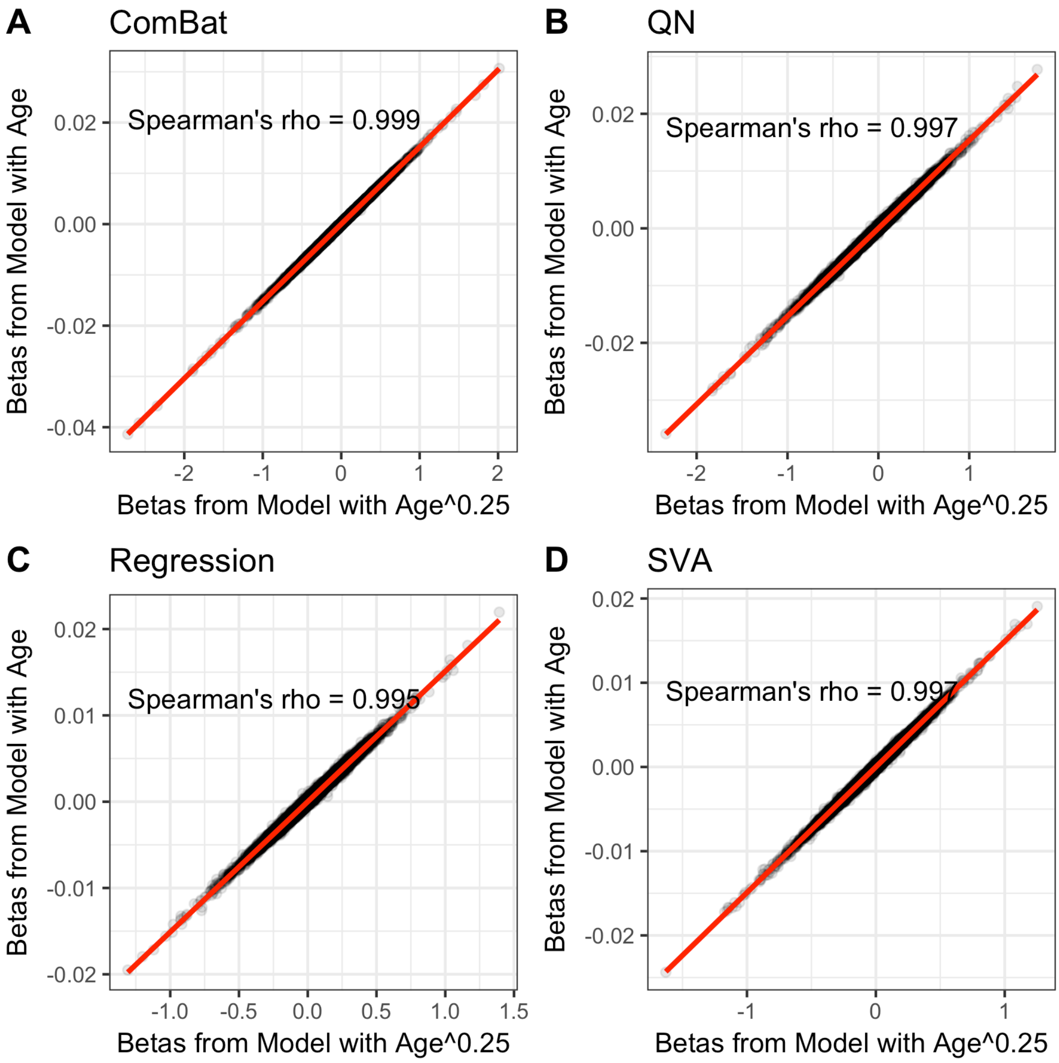


**Supplemental Figure 4.** Scatter plots of the dependency between beta1 coefficients from the linear regression models that explain gene expression using age in comparison to age$^{0.25}$. Gene expression was corrected using ComBat **(A)**, QN only **(B)**, regression **(C)** and SVA **(D)**. Linear regression line between the variables (in red) and Spearman correlation estimate is shown on the graph.

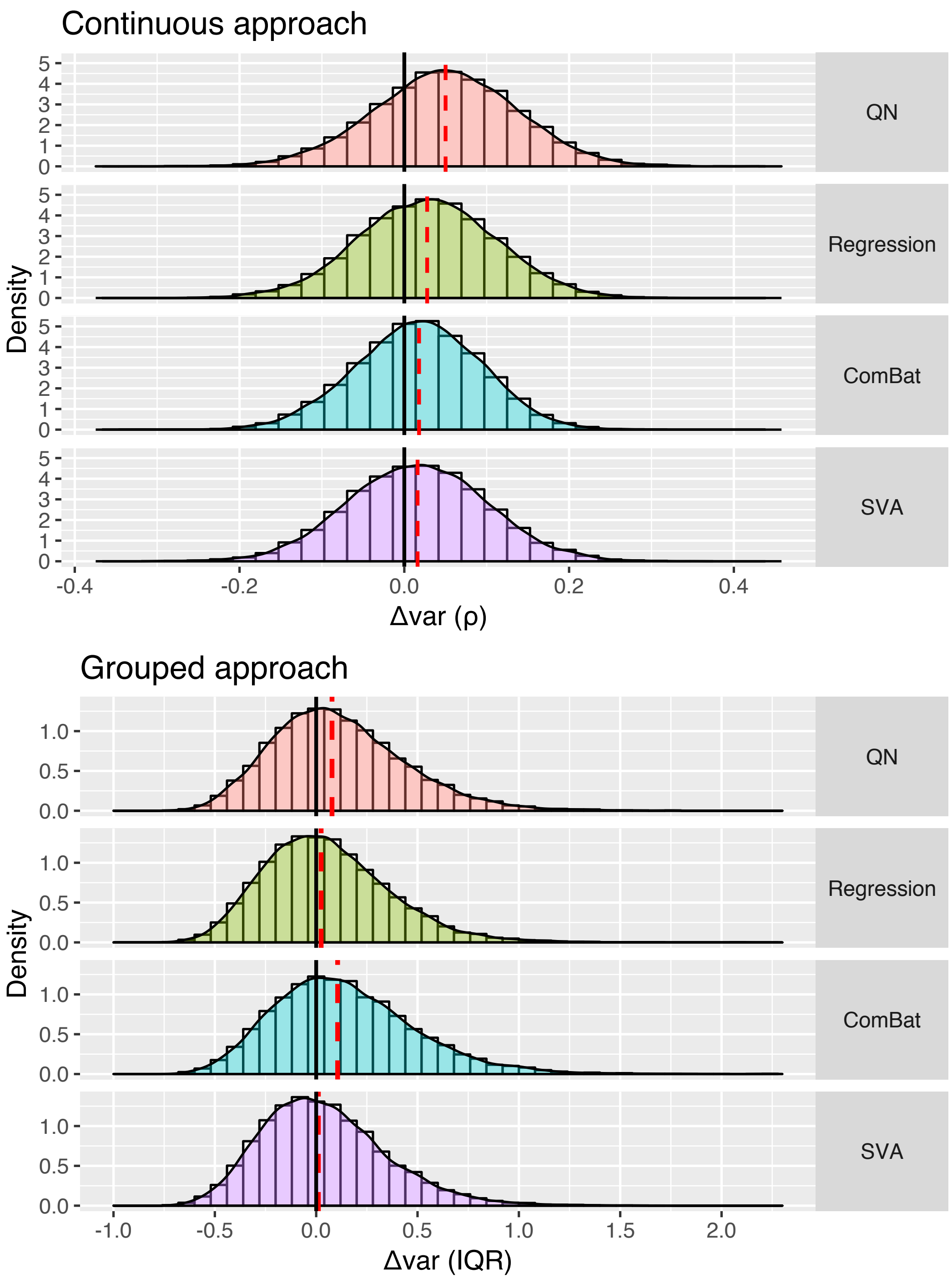


**Supplemental Figure 5.** Comparison of the distributions of the differential variability measures, Δvar (ρ) and Δvar (IQR), obtained in the expression data corrected with QN, Regression, SVA and ComBat. The black straight line depicts zero, the red dashed line – median of the distribution.

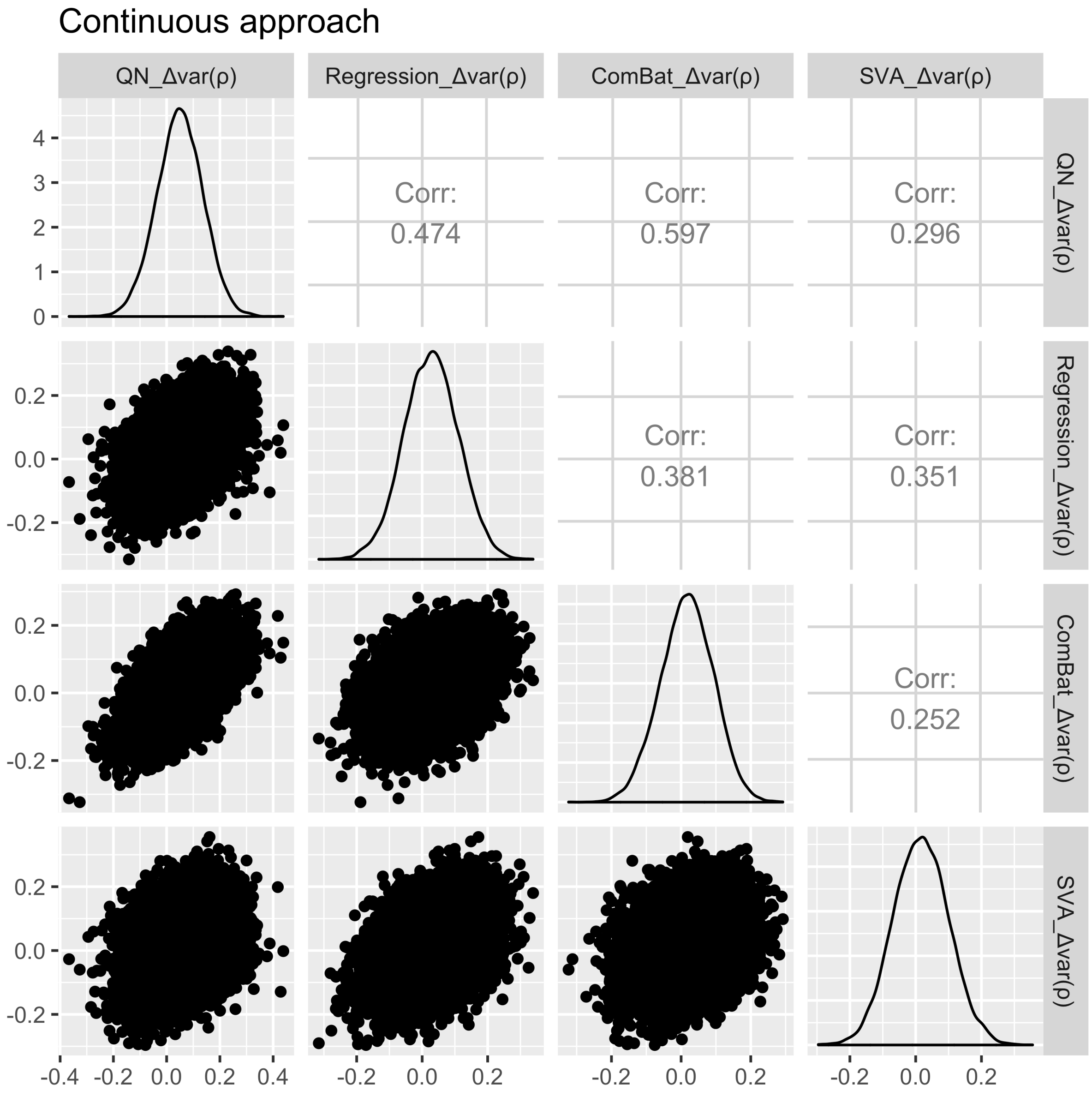


**Supplemental Figure 6.** Combination of comparisons of the variability measures (Δvar) obtained in the continuous approach between different ways of data pre-processing: correction with quantile normalisation only, linear regression, SVA and ComBat. Scatter plots between the differential variability measures obtained after different corrections are displayed in the lower triangle of the matrix, distributions of the differential variability measures are located on the diagonal and Spearman correlation estimates between the variability measures from different corrections are shown in the upper triangle of the matrix.

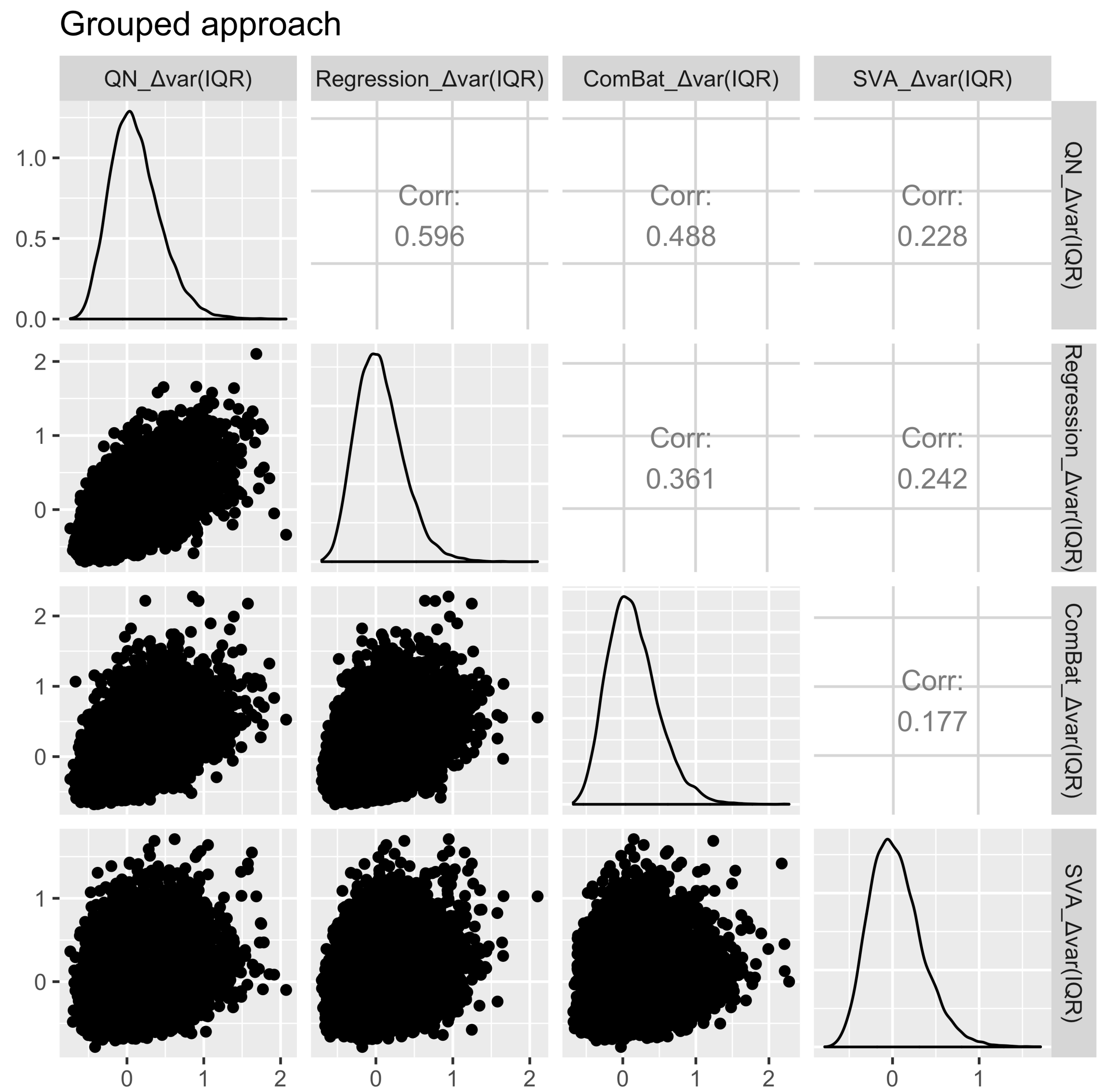


**Supplemental Figure 7.** Combination of comparisons of the variability measures (Δvar) obtained in the grouped approach between different ways of data pre-processing: correction with quantile normalisation only, linear regression, SVA and ComBat. Scatter plots between the differential variability measures obtained after different corrections are displayed in the lower triangle of the matrix, distributions of the differential variability measures are located on the diagonal and Spearman correlation estimates between the differential variability measures from different corrections are shown in the upper triangle of the matrix.

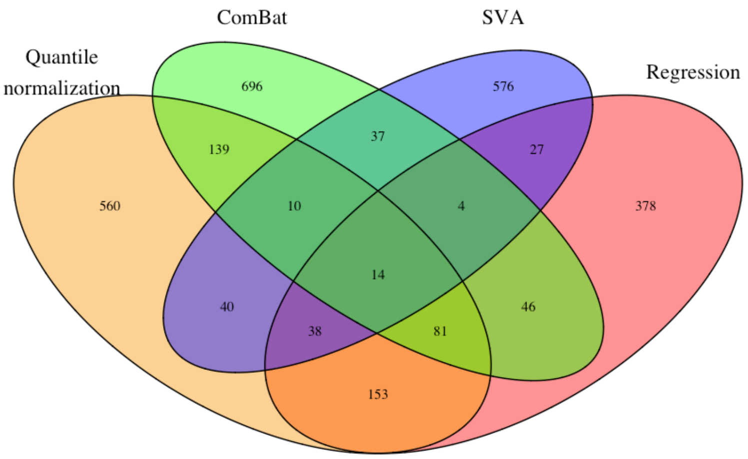


**Supplemental Figure 8.** Venn diagram of the differentially variable genes found in the grouped approach when expression data was corrected with QN, linear regression, SVA and ComBat.

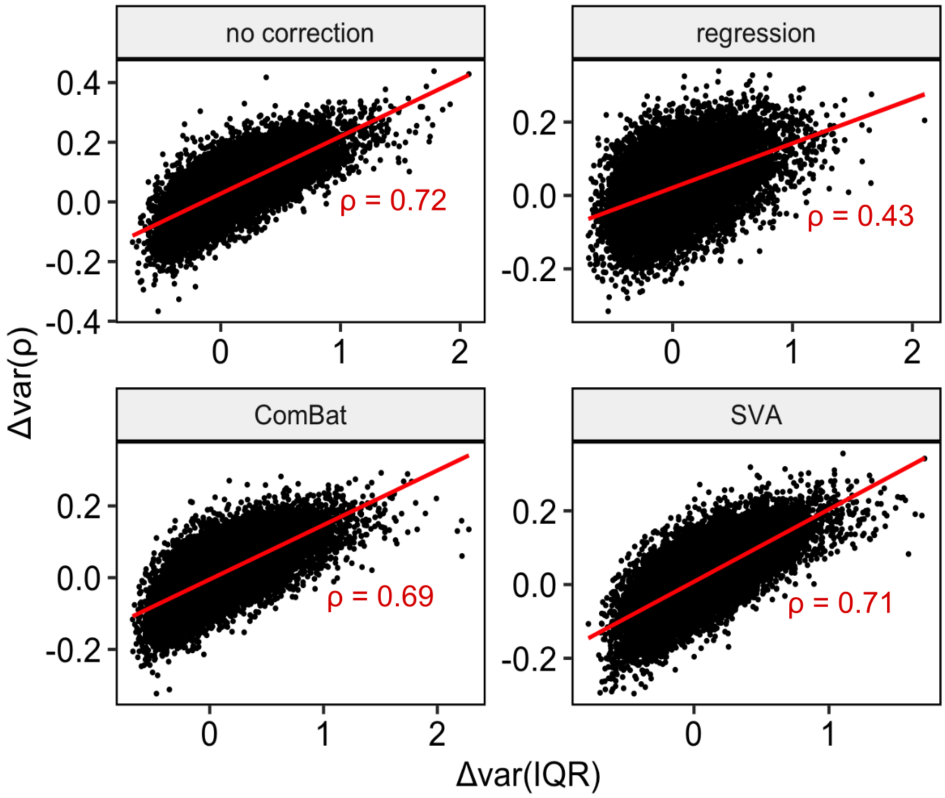


**Supplementary Figure 9.** Scatter plot of the relationship between the change in the variability measures for the continuous and grouped approaches in data, corrected with QN, ComBat, regression and SVA corrections. Linear regression line between the variables (in red) and Spearman correlation estimate are shown on the graph.

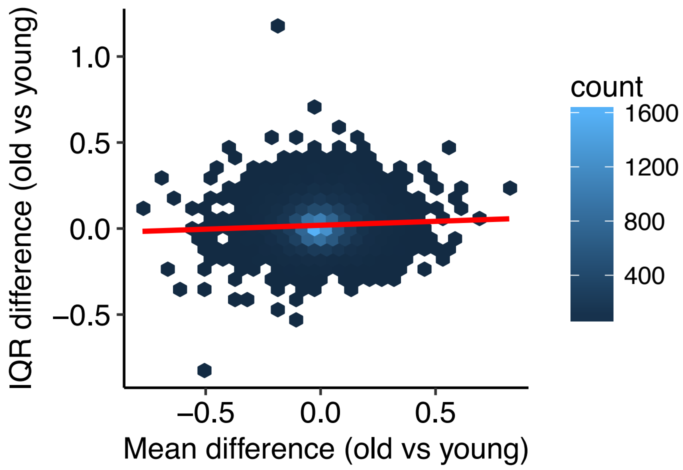


**Supplemental Figure 10.** Relationship between the change in gene expression level and variability: x axis depicts difference between mean of the old group and mean of the young group distribution (generated by resampling young group with sample size of 22 for 10,000 times and calculating the mean), y axis illustrates difference between interquartile range (IQR) for the old and mean IQR from the young distribution (generated by resampling young group with sample size of 22 for 10,000 times and calculating IQR).

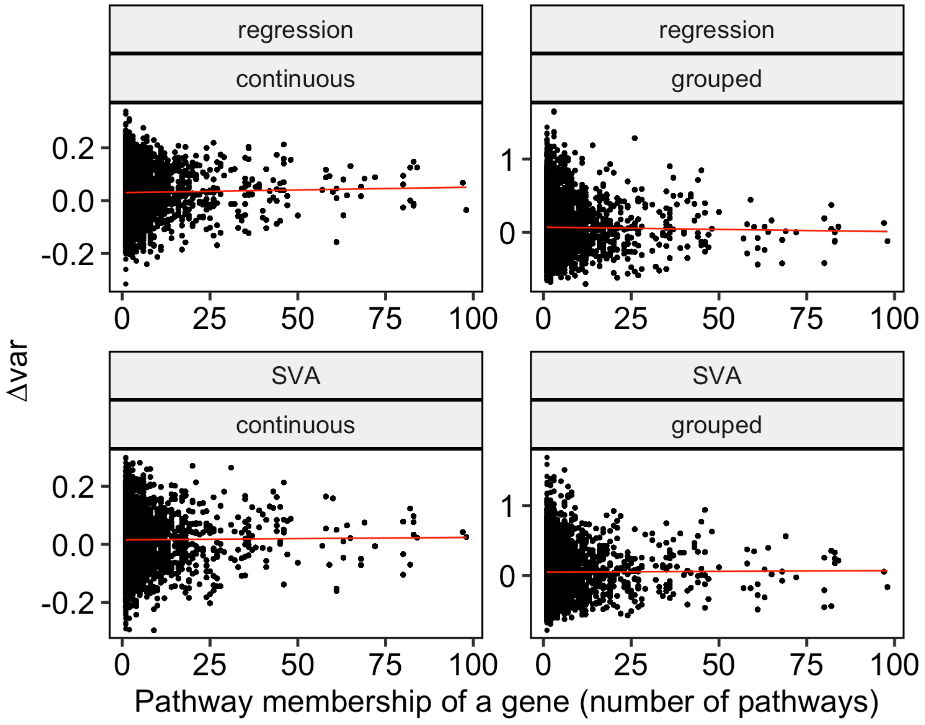


**Supplemental Figure 11.** Scatter plots that illustrate relationship between the pathway membership of a gene and its change in variability measure for the continuous and grouped approaches in data corrected with regression and SVA. Linear regression line between the variables is shown red on the graph.
